# Genomic analyses underpin the feasibility of concomitant genetic improvement of milk yield and mastitis resistance in dairy sheep

**DOI:** 10.1101/577015

**Authors:** Georgios Banos, Emily L. Clark, Stephen J. Bush, Prasun Dutta, Georgios Bramis, Georgios Arsenos, David A. Hume, Androniki Psifidi

**Affiliations:** Scotland’s Rural College, Edinburgh, Easter Bush, Midlothian, EH25 9RG, UK; The Roslin Institute, University of Edinburgh, Easter Bush, Midlothian, EH25 9RG, UK; School of Veterinary Medicine, Aristotle University of Thessaloniki, Thessaloniki, 54124, Greece; Nuffield Department of Clinical Medicine, University of Oxford, John Radcliffe Hospital, Headington, Oxford, OX3 9DU, UK; Mater Research Institute-University of Queensland, Translational Research Institute, Woolloongabba, Qld 4102, Australia; Royal Veterinary College, University of London, Hatfield, AL9 7TA, UK

**Author notes:** Correspondence: Dr Androniki Psifidi.

## Abstract

Milk yield is the most important dairy sheep trait and constitutes the key genetic improvement goal via selective breeding. Mastitis is one of the most prevalent diseases, significantly impacting on animal welfare, milk yield and quality, while incurring substantial costs. Our objectives were to determine the feasibility of a concomitant genetic improvement programme for enhanced milk production and resistance to mastitis. Individual records for milk yield and four mastitis-related traits were collected monthly throughout lactation for 609 ewes of the Chios breed. All ewes were genotyped with a mastitis specific custom-made 960 single nucleotide polymorphism array. We performed genomic association studies, (co)variance component estimation and pathway enrichment analysis, and characterised gene expression levels and the extent of allelic expression imbalance. Presence of heritable variation for milk yield was confirmed. There was no significant genetic correlation between milk yield and mastitis. Environmental factors appeared to favour both milk production and udder health. Four Quantitative Trait Loci (QTLs) affecting milk yield were detected on chromosomes 2, 12, 16 and 19, in locations distinct from those previously identified to affect mastitis resistance. Pathways, networks and functional gene clusters for milk yield were identified. Seven genes (*DNAJA1, DNAJC10, FGF10, GHR, HMGCS1, LYPLA1, OXCT1*) located within the QTL regions were highly expressed in both the mammary gland and milk transcriptome, suggesting involvement in milk synthesis and production. Furthermore, the expression of four genes (*DNAJC10, FGF10, OXCT1, EMB*) was enriched in immune tissues implying a favourable pleiotropic effect or likely role in milk production during udder infection. In conclusion, the absence of genetic antagonism between milk yield and mastitis resistance suggests that simultaneous genetic improvement of both traits be achievable. The detection of milk yield QTLs with the mastitis array underpins the latter’s utility as a breeding tool for the genetic enhancement of both traits.

## Introduction

The world’s commercial dairy sheep industry is primarily concentrated in Mediterranean countries and linked to local breeds; milk is mostly used to produce high quality cheeses and other dairy products. Milk yield represents more than two thirds of the total income of the dairy sheep sector [1] and, therefore, increasing milk yield is the most important and sometimes only objective of selective breeding. Milk production traits in dairy sheep are moderately to highly heritable [2, 3] and amenable to improvement with traditional selective breeding programmes based on pedigree and phenotypic data. Indeed, such programmes have been established in many sheep populations over recent decades [2, 4]. Incorporation of genomic information in some breeding programmes (e.g. French Lacaune, Spanish Churra, Italian Sarda) has led to an acceleration of the genetic improvement outcomes.

The Greek Chios breed is considered to be among the most productive and prolific dairy sheep breeds worldwide [5]. A traditional breeding programme for the enhancement of milk yield has been in place since year 2000 for this breed, leading to substantial improvement in this trait. However, further increases in milk yield may be achieved with the use of relevant genomic information.

Beyond simply increasing milk production, the dairy sheep industry faces challenges such as the need to offer healthy products to consumers, addressing animal welfare, and ensuring the long-term competitiveness and sustainability of the sector. Mastitis is the most prevalent and costly disease in the dairy industry due to reduced and discarded milk, early involuntary culling of animals, and veterinary services and labour costs [6, 7]. The disease also poses a potential threat of zoonosis and antimicrobial resistance if antibiotic treatment is not applied carefully [6–8]. Moreover, mastitis is a welfare concern because of associated pain, anxiety and restlessness, and upsets the normal feeding behaviour of the animals [9]. Host resistance to mastitis is a moderately heritable trait [7]. Recently, an ovine custom made mastitis specific 960-SNP DNA array was built to facilitate genetic selection and improvement of animal resistance to mastitis in dairy sheep [10] [11] [12] [13]. We previously used this array in a genomic association study and detected five quantitative trait loci (QTLs) for mastitis resistance in Chios sheep [10].

In the present study, we examined the genetic and genomic relationship between milk yield and mastitis resistance in the Chios sheep, using pedigree and genomic information. The relationship between the two traits is crucial if mastitis resistance is to be included in selective breeding goals together with increased milk yield. We estimated genetic parameters and investigated whether relevant QTLs for milk yield exist in previously identified mastitis-specific genomic regions. We also performed pathway analysis and examined gene expression and allelic expression imbalance to characterise the genes located under the QTL regions in relation with milk yield and mastitis resistance.

## Materials and Methods

### Ethical statement

The study was approved by the Ethics and Research Committee of the Faculty of Veterinary Medicine, Aristotle University of Thessaloniki, Greece. Permits for access and use of the commercial farms were granted by the farm owners, who were members of the Chios Sheep Breeders’ Cooperative “Macedonia”. During sampling, animals were handled by qualified veterinarians. Permission to qualified veterinarians to perform milk and blood sampling was granted by the National (Greek) Legislature for the Veterinary Profession, No. 344/29-12-2000.

### Animals, sampling and phenotyping

Animals used in the present study included 609 purebred Chios dairy ewes raised in four commercial farms in Greece. Complete pedigree data were available for these animals. Ewes were in their first or second lactation. Daily milk yield was recorded on each animal on the day of monthly visits to the farms during the first five months of lactation. The first milk yield record was obtained at least three days after lamb weaning (ca. 42 days post lambing). Total number of individual animal records collected amounted to 2,436. Animal records for clinical mastitis occurrence and three mastitis indicator traits (milk somatic cell count, California Mastitis Test score and total viable bacterial count in milk) were also collected at the time of these visits by a qualified veterinarian, as described previously [10]. The three mastitis indicator traits may capture subclinical mastitis incidences and reflect the general health status of the udder. Peripheral blood samples were taken from each ewe in 9 ml K_2_EDTA Vacutainer blood collection tubes (BD diagnostics) by jugular venepuncture for genomic DNA extraction.

### Genetic parameter estimation

Genetic parameters for milk yield were estimated using the following basic mixed model:

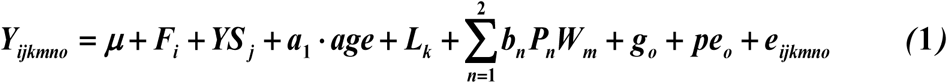

Where: Y = record of ewe *o* in week of lactation *m*

*µ* = overall mean

*F* = fixed effect of flock (farm) *i*

*YS* = fixed effect of year-season of lambing *j*

*α*_1_ = linear regression on age at lambing (*age*)

*L* = fixed effect of lactation number *k*

*W* = fixed effect of week of lactation (i.e. week post-lambing) *m*

*b*_*n*_ = fixed regression coefficient on week of lactation *m* (order n=2)

*P_n_* = orthogonal polynomial of week *m* (order n=2)

*g* = random additive genetic effect of ewe *o*, including pedigree genetic relationships among animals

*pe* = random permanent environment effect of ewe *o*

*e* = random residual effect

Heritability and repeatability estimates were derived from the variance components calculated for the random effects in model (1). In a separate analysis, the additive genetic and permanent environment effects in model (1) were replaced by interactions of the latter with second-order polynomial functions of week of lactation. The choice of polynomial order was decided after testing sequentially increasing orders with the log-likelihood test. This analysis resulted in distinct variance component and genetic parameter estimates by week of lactation, which were then combined to derive average heritability and repeatability estimates for early (weeks of lactation 1-7), mid (weeks 8-17) and late (weeks 18-24) lactation. In addition, genetic correlations between milk yields measured at different lactation stages were calculated based on corresponding genetic covariance estimates. A smoothed lactation curve adjusted for all fixed effects in the model was also derived.

Finally, bivariate analyses of milk yield and each one of the four mastitis related traits were conducted using model (1); outcomes from these analyses were used to estimate phenotypic and genetic correlations between traits.

All statistical analyses in the present study were conducted with ASReml v4.0 [14].

### Genomic association studies

DNA was extracted from blood buffy coat as described previously [15].

All animals were genotyped with a customised mastitis specific 960 SNP DNA array containing SNPs located on chromosomes 2, 3, 5, 12, 16 and 19. Briefly, this array was built based on QTLs for mastitis resistance found to segregate in multiple different dairy sheep breeds. For the design of this custom-made array, SNPs were selected from 50K and 800K SNP ovine DNA arrays, as well as available re-sequencing data. The average density of the array was 1 SNP every 23 Kb (for more details see [10]). This genomic tool was built within an FP7 European research project (http://cordis.europa.eu/result/rcn/163471_en.html). Genotypes at each SNP locus were subjected to quality control measures using the following thresholds: minor allele frequency >0.05, call rate >95% and Hardy-Weinberg equilibrium P>10^−4^. After quality control, 710 SNP markers remained for further analysis.

Possible population stratification was investigated with the use of the genomic relationship matrix among individual animals. This matrix was converted to a distance matrix that was then used to conduct multidimensional scaling analysis using the GenABEL package of R [16].

Individual ewe phenotypes were residuals resulted after fitting a model that included all fixed effects of model (1); thus, phenotypic records were adjusted for all these environmental effects. Separate phenotypes were derived for the entire lactation (overall) and for each lactation stage (early, mid, late) as described above. In all cases, GEMMA v0.94.1 [17] was used to conduct genomic association analyses based on a mixed model that included the genomic relationship matrix among individual ewes as a polygenic effect. After Bonferroni correction for multiple testing, the significance threshold for nominal P=0.05 was set at P=7.04×10^−5^ and a suggestive threshold (accounting for one false positive per genome scan) was set at P=1.41×10^−3^.

Statistically significant SNPs from the genomic association analyses were further examined with a mixed model that included the fixed effects of model (1), the fixed effect of the SNP genotype and the random effect of the animal including the pedigree relationship matrix. Additive (a) and dominance (d) effects, and the proportion of additive genetic variance due to each SNP locus (pVA) were calculated as follows:

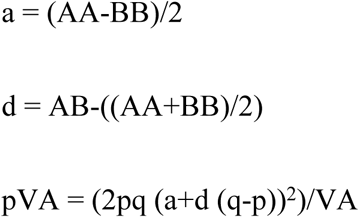

where AA, BB and AB were the marginal means of the respective genotype, p and q the corresponding frequencies of alleles A and B at the SNP locus, and VA the estimated additive genetic variance, derived with model (1). All analyses were conducted with ASReml v4.0 [14].

Linkage disequilibrium (LD) among significant SNPs was calculated based on the *r^2^* value using PLINK v1.9 [18]. Blocks of LD in regions harbouring significant SNPs were visualised using Haploview v4.2 [19].

All significant (post-Bonferroni correction) and suggestive SNPs identified in the genomic analysis for milk yield were mapped to the reference genome and annotated using the variant effect predictor (http://www.ensembl.org/Tools/VEP) tool within the Ensembl database and the Oar v3.1 assembly. Moreover, genes located around (0.5 Mb windows upstream and downstream) the significant markers -in the candidate regions for milk yield-were annotated using the BioMart data mining tool (http://www.ensembl.org/biomart/martview/) and the Oar v3.1 assembly.

### Pathway and functional enrichment analysis

The list of annotated genes located within the QTL regions for milk yield were analysed with the Ingenuity Pathway Analysis (IPA) programme (www.ingenuity.com) in order to identify canonical pathways and gene networks constructed by the products of these genes. IPA constructs multiple possible upstream regulators, pathways and networks which may be associated with the biological mechanism underlying the studied trait. The analysis is based on data from large-scale causal networks derived from the Ingenuity Knowledge Base. IPA then infers the most suitable pathways and networks based on their statistical significance, after correcting for a baseline threshold [20]. The IPA score in the constructed networks can be used to rank these networks based on the P-values obtained using Fisher’s exact test (IPA score or P*-*score = –log_10_(P value)).

The list of candidate genes was also analysed against an *Ovis aries* background using the Database for Annotation, Visualization and Integrated Discovery (DAVID) [21] to examine gene set enrichment. We determined the corresponding gene ontology terms and performed functional annotation clustering analysis to detect gene enrichment. The enrichment score calculated by the DAVID software package is a modified Fisher’s exact test P*-*value; an enrichment score greater than unity reflects over-representation of the respective functional category.

### Gene expression analysis

Genes contributing to milk production are likely to be expressed in milk somatic cells, mammary gland, and other organs such as the liver and kidney that provide nutrients and regulate the electrolytes needed for lactosynthesis and the production of milk. We also reasoned that the expression of genes with pleiotropic effects would be associated with both milk yield and resistance to mastitis, and/or expressed in both mammary gland and immune related tissues. To assess the expression profiles of genes located in the candidate regions for milk yield, we obtained publicly available data from an RNA-seq characterisation of the milk transcriptome of two Spanish dairy sheep breeds, Churra and Assaf, where milk somatic cells of eight individual sheep (four from each breed) had been sampled throughout lactation at 10, 50, 120 and 150 days after lambing [22, 23]. Individual milk yield and milk somatic cell count records were also available for the sheep used in the latter study [23]. To supplement this data, we used publicly available RNA-Seq data from a high-resolution atlas of gene expression across tissues and cell types from all major organ systems in sheep [24, 25]. The sheep gene expression atlas, which includes 437 RNA-Seq libraries was produced using six Texel × Scottish Blackface sheep [24]. An additional 83 RNA-Seq libraries from a Texel trio (ewe, lamb and ram) were included in the sheep gene expression atlas [25]. We extracted data pertaining to the mammary glands, liver and kidneys. Since we were interested in detecting genes related to both milk yield and mastitis, we also extracted the expression level of the genes under consideration in immune-related tissues, specifically hemolymph nodes, mesenteric, popliteal, prescapular and submandibular lymph nodes, peripheral blood mononuclear cells, blood leukocytes, monocyte-derived macrophages, bone marrow derived macrophages, alveolar macrophages, and tonsils.

Expression levels for all samples, were estimated using Kallisto v0.42.4 [26]. Expression was reported for each protein-coding transcript as the number of transcripts per million, and then summarised to the gene-level (as in [27]). Heatmaps were drawn using the heatmap.2 function of the R package gplots v3.0.1, in order to demonstrate expression enrichment in the different tissues and lactation stages.

The relationship of the expression level of each gene in the milk transcriptome with milk yield and milk somatic cell count was assessed in the Spanish sheep data using the following linear model:

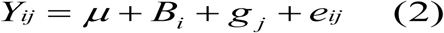

where Y = record of ewe (milk yield or milk somatic cell count), *µ* = overall mean, *B* = fixed effect of breed *i*, *g* = fixed effect of the mean expression of gene *j*, *e* = random residual effect.

The nominal significance threshold in this analysis was set at P=0.05. Since genes were located within four QTL regions, an FDR adjustment for multiple testing was applied, setting the significance threshold at P=0.0167. These analyses were conducted with ASReml v4.0 [14].

To identify significant expression differences amongst genes located in the milk yield candidate regions in sheep with low, medium and high milk somatic cell count we performed Tukey’s Test using the statistical package R v3.0.1.

### Variant calling and allelic expression imbalance analysis

Much of the genetic variation in genes that control a quantitative trait is likely to affect their transcriptional regulation. In fact, many quantitative traits associated with altered gene expression, and trait-associated loci are enriched for eQTLs (Nicolae et al., 2010). If an individual is heterozygous for a *cis*-acting mutation it is expected that the two alleles of the gene will be expressed unequally causing allelic expression imbalance. Measuring the relative expression levels of two alleles using RNA-Seq may lead to the identification of *cis*-acting SNPs or haplotypes [28–31]. To identify any *cis*-QTLs affecting the genes located in the candidate regions for milk yield we obtained the raw RNA-Seq data for mammary gland tissue from three adult female Texel × Scottish Blackface sheep from the sheep gene expression atlas [24]. The aligner HISAT2 (v2.0.4) [32], was used to produce the BAM files as previously described [24]. Variants were called using BCFtools [33] mpileup (v1.4) with parameters --max-depth 1000000 --min-MQ 60, followed by BCFtools call (v1.4) with parameters -m (allow multiallelic variants) and -v (variant only). The minimum MAPQ (mapping quality) score was chosen to focus on uniquely mapped reads for variant calling. The resulting VCF file contained both SNPs and indels. The exonic variants of the protein coding genes located in the milk yield candidate regions were obtained from each VCF file using the program GTF_Extract (v0.9.1) (https://github.com/fls-bioinformatics-core/GFFUtils/blob/master/docs/GTF_extract.rst) and BEDtools [34] intersect (v2.25.0) based on gene annotations from Ovis_aries.Oar_v3.1. The putative functional impact of each variant on the encoded proteins was predicted using SnpEff v4.3 [35] with the parameter – onlyProtein (only annotate protein-coding variants). BCFtools norm (v1.4) with parameter – d was used to remove duplicated VCF records that arose due to duplicated exon coordinates in the GTF file (that is, exons present in more than one transcript). Finally, VCFs from each animal were filtered to obtain only biallelic heterozygous SNPs, using BCFtools ‘view’ (v1.4). For the allelic expression imbalance analysis we focused on biallelic heterozygous exonic SNPs, since the non-exonic variants may signify transcriptional noise in mRNA sequencing and contribute potential errors in the analysis.

Read counts for both the reference and alternate allele were obtained using allelecounter v0.6 (https://github.com/secastel/allelecounter) with parameters --min_cov 4, --min_baseq 20 and --min_mapq 60 and --max_depth 10000. Allelic expression imbalance, per gene, was estimated using MBASED (Meta-analysis Based Allele-Specific Expression Detection) [36] with parameters isPhased=FALSE, numSim=10^6, BPPARAM=SerialParam(). MBASED allelic expression imbalance estimates were derived by combining information across individual heterozygous SNP within a gene. Only variants with >10 reads in either reference or alternate allele were used. We retained only those genes with Benjamin-Hochberg [37] adjusted P-value ≤0.05 and major allele frequency ≥ 0.7.

## Results

### Descriptive statistics

An average daily milk yield of 1,912 grams (g) was produced in the studied sheep population with a standard deviation of 713 g, a maximum of 4,597 g and a minimum of 210 g. As expected, milk yield decreased as lactation progressed [38].

### Genetic parameters

Estimates of heritability and repeatability of milk yield (Table 1) were derived for the entire lactation as well as different stages of lactation defined as early, mid and late. Statistically significant (*P<*0.05) moderate trait heritabilities (0.19-0.28) and repeatabilities (0.69-0.76) were estimated across all lactation stages. Moreover, the genetic correlations between milk yield in different lactation stages were significantly (*P*<0.05) positive. However, the genetic correlation between early and late lactation was moderate (0.60) and significantly less than one. In practical terms, lactation onset, peak lactation and lactation persistence may have partly separate genetic control. Genetic correlations between milk and mastitis traits were not significantly different from zero (*P*>0.05). Negative phenotypic correlations were observed between these traits (*P*<0.05), indicative of favourable environmental effects to both production and health (Table 2).

**Table 1.** Heritability (h^2^) and repeatability (r) estimates of daily milk yield in Chios sheep by lactation stage and across the entire lactation; standard errors in parentheses.

| Parameter | Early lactation<br>(1-7 weeks) | Mid lactation<br>(8-17 weeks) | Late lactation<br>(18-24 weeks) | Overall<br>lactation |
| --- | --- | --- | --- | --- |
| $h^2$ | 0.28 (0.06) | 0.19 (0.06) | 0.23 (0.06) | 0.23 (0.06) |
| $r$ | 0.76 (0.02) | 0.69 (0.02) | 0.71 (0.02) | 0.71 (0.02) |

**Table 2.** Estimates of phenotypic and genetic correlations between milk yield and four mastitis traits in Chios sheep; standard errors in parentheses.

| **Mastitis trait** | **Phenotypic correlation** | **Genetic correlation** |
| --- | --- | --- |
| **SCC** | -0.18 (0.04)* | -0.12 (0.14) |
| **CMT** | -0.18 (0.04)* | -0.12 (0.13) |
| **TVC** | -0.10 (0.03)* | -0.11 (0.14) |
| **CM** | -0.07 (0.04) | -0.09 (0.19) |
| SCC: milk somatic cell count, CMT: California Mastitis Test score, TVC: total bacterial count in milk, CM: clinical mastitis occurrence; *Significantly different from zero ($P < 0.05$) | | |

### Genomic association studies

Separate genomic association analyses were conducted for milk yield in early, mid, late and overall lactation. Multidimensional scaling analysis of the studied population revealed no substructure. In general, similar genomic associations were detected for milk yield in middle, late and overall lactation but distinct associations were observed in early lactation. We identified a genome-wide significant association after Bonferroni correction for multiple testing on chromosome 19 (*P*= 1.28 × 10^−5^) and three suggestive associations on chromosomes 2 (*P*= 4.30 × 10^−4^), 12 (*P*= 3.65 × 10^−4^) and 16 (*P*= 6.07 × 10^−4^). Details of SNPs associated with milk yield are shown in Table 3. Manhattan plots and corresponding Q-Q plots displaying genomic association results are shown in Fig. 1 and Fig. 2, respectively.

**Table 3.** List of Single Nucleotide Polymorphisms (SNPs) associated with milk yield in Chios sheep.

| Lactation stage | SNP | Chr (position) | P-value | Add(P-value) | Dom(P-value) | pVA | p | q |
| --- | --- | --- | --- | --- | --- | --- | --- | --- |
| **Early 1-7 weeks** | OAR12_23075585 | 12(20050780) | 3.35E-04 | 0.07(0.09) | 0.07(0.10) | 0.01 | 0.62 | 0.38 |
|  | oar3_OAR12_19689222 | 12(19689222) | 3.65E-04 | 0.05(0.36) | 0.12(0.03) | 0.01 | 0.73 | 0.27 |
|  | oar3_OAR12_19269103 | 12(19269103) | 4.66E-04 | -0.03(0.33) | 0.05(0.02) | 0.01 | 0.73 | 0.27 |
|  | oar3_OAR12_19500329 | 12(19500329) | 6.98E-04 | 0.08(0.02) | 0.02(0.64) | 0.02 | 0.6 | 0.4 |
|  | oar3_OAR12_19624437 | 12(19624437) | 6.77E-04 | 0.05(0.30) | 0.09(0.10) | 0.01 | 0.72 | 0.28 |
|  | oar3_OAR12_19840123 | 12(19840123) | 9.80E-04 | 0.04(0.30) | 0.08(0.10) | 0.01 | 0.68 | 0.32 |
|  | oar3_OAR16_33078067 | 16(33078067) | 6.07E-04 | -0.09(0.00) | 0.20(0.05) | 0.05 | 0.96 | 0.04 |
| **Middle 8-17 weeks** | OAR19_25259444 | 19(23804520) | 7.03E-05 | -0.15 (0.00) | 0.05 (0.25) | 0.16 | 0.48 | 0.52 |
|  | oar3_OAR19_24119431 | 19(24119431) | 9.03E-04 | -0.14(0.00) | -0.02(0.53) | 0.15 | 0.48 | 0.52 |
|  | OAR19_25513179 | 19(24010793) | 1.70E-03 | -0.15(0.00) | -0.01(0.77) | 0.16 | 0.58 | 0.42 |
|  | OAR16_34906481 | 16(32156238) | 1.35E-03 | -0.08(0.02) | 0.07(0.05) | 0.05 | 0.90 | 0.10 |
|  | OAR2_133418483 | 2(125230366) | 1.48E-03 | 0.08(0.51) | -0.10(0.44) | 0.06 | 0.92 | 0.08 |
|  | OAR2_133088440 | 2(124907852) | 2.28E-04 | 0.23(0.00) | 0.19(0.03) | 0.05 | 0.82 | 0.18 |
|  | oar3_OAR2_124936445 | 2(124936445) | 1.11E-03 | 0.12(0.06) | 0.07(0.30) | 0.03 | 0.78 | 0.22 |
| **Late 18-24 weeks** | OAR19_25259444 | 19(23804520) | 1.41E-04 | -0.15(0.00) | -0.02(0.57) | 0.14 | 0.48 | 0.52 |
|  | oar3_OAR19_24745933 | 19(24745933) | 2.24E-04 | 0.10(0.00) | -0.10(0.00) | 0.07 | 0.54 | 0.46 |
|  | OAR19_25830151 | 19(24342061) | 1.35E-03 | 0.07(0.07) | -0.00(0.87) | 0.02 | 0.72 | 0.28 |
|  | oar3_OAR19_24707843 | 19(24707843) | 7.91E-04 | -0.13(0.00) | -0.09(0.03) | 0.06 | 0.64 | 0.36 |
|  | oar3_OAR19_23656789 | 19(23656789) | 6.29E-04 | -0.11(0.00) | -0.06(0.11) | 0.07 | 0.52 | 0.48 |
|  | oar3_OAR2_124936445 | 2(125000000) | 5.76E-04 | 0.11(0.02) | -0.00(0.93) | 0.05 | 0.78 | 0.22 |
|  | OAR2_133088440 | 2(124907852) | 4.30E-04 | 0.19(0.00) | 0.09(0.18) | 0.06 | 0.82 | 0.18 |
| **Overall** | OAR19_25259444 | 19(23804520) | 1.28E-05 | -0.14(0.00) | -0.00(0.84) | 0.12 | 0.48 | 0.52 |
|  | oar3_OAR19_24032312 | 19(24032312) | 2.70E-04 | -0.11(0.00) | -0.08(0.03) | 0.05 | 0.6 | 0.4 |
|  | oar3_OAR19_24707843 | 19(24707843) | 2.90E-04 | -0.12(0.00) | -0.07(0.04) | 0.06 | 0.64 | 0.36 |
|  | OAR19_25513179 | 19(24010793) | 4.16E-04 | -0.11(0.00) | -0.06(0.07) | 0.06 | 0.58 | 0.42 |
|  | oar3_OAR19_24119431 | 19(24119431) | 4.90E-04 | -0.11(0.00) | -0.09(0.01) | 0.08 | 0.48 | 0.52 |
|  | oar3_OAR19_23929524 | 19(23929524) | 5.62E-04 | 0.11(0.00) | -0.03(0.29) | 0.07 | 0.48 | 0.52 |
|  | oar3_OAR19_24745933 | 19(24745933) | 9.80E-04 | 0.10(0.00) | -0.05(0.15) | 0.06 | 0.54 | 0.46 |
|  | oar3_OAR19_23891277 | 19(23891277) | 1.17E-03 | -0.10(0.00) | -0.05(0.13) | 0.05 | 0.54 | 0.46 |
|  | oar3_OAR19_23656789 | 19(23656789) | 1.46E-03 | -0.10(0.00) | -0.05(0.15) | 0.06 | 0.52 | 0.48 |
|  | OAR2_133088440 | 2(124907852) | 9.55E-04 | 0.20(0.00) | 0.12(0.06) | 0.05 | 0.82 | 0.18 |
|  | OAR2_133418483 | 2(125230366) | 1.47E-03 | 0.10(0.30) | -0.08(0.43) | 0.05 | 0.92 | 0.08 |

P-value: P-value from genomic association study; additive allele substitution effect (ADD) and corresponding P-value; dominance effect (DOM) and corresponding P-value; pVA: proportion of the genetic variance explained by the SNP; p and q allelic frequencies

**Fig 1.**
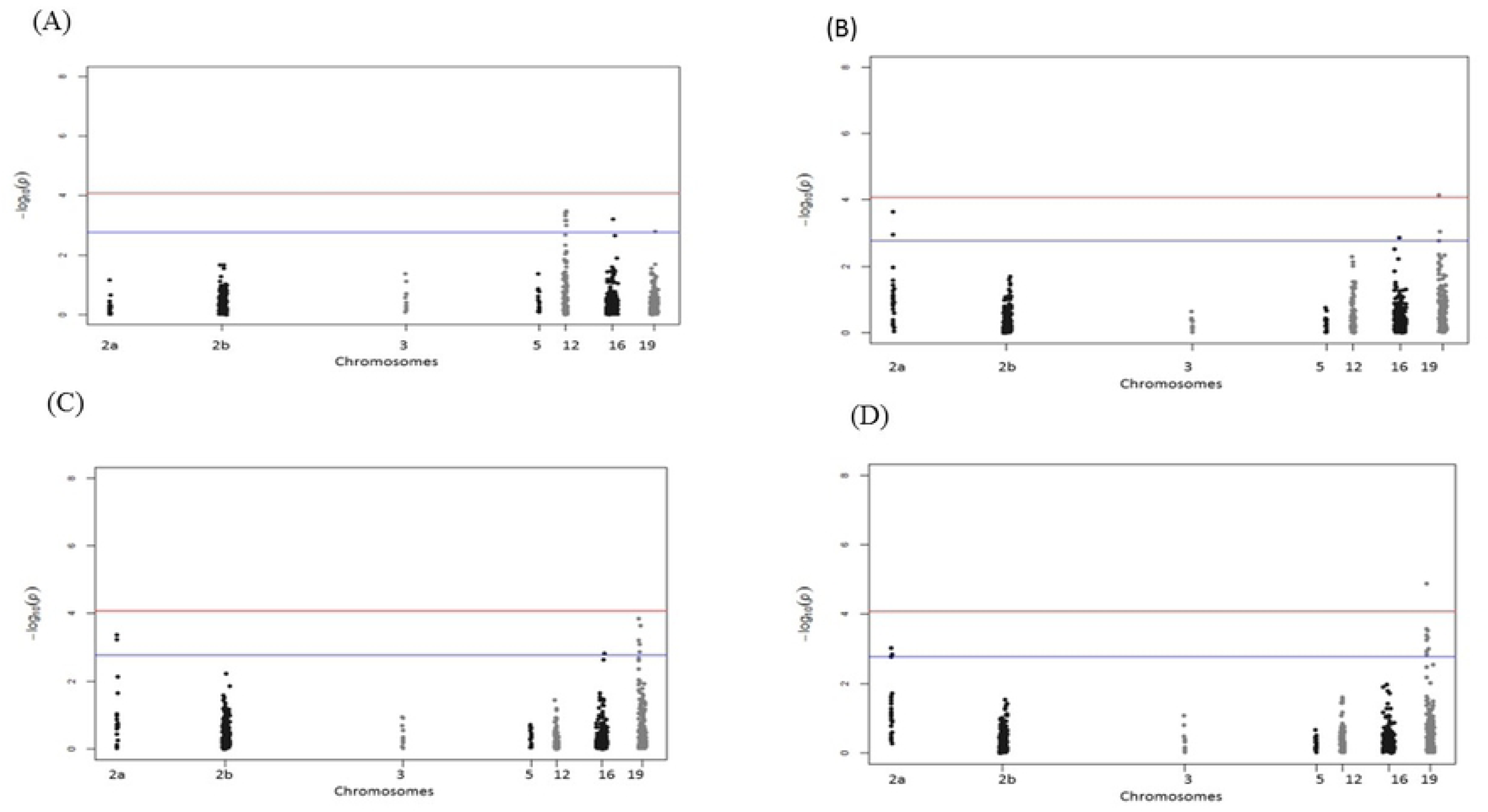
Manhattan plots displaying the genomic association results for milk yield in Chios sheep. Manhattan plots for milk yield in early (A), mid (B), late (C), and overall (D) lactation. Genomic location is plotted against −log_10_(*P*). Red and blue lines, respectively, are thresholds for significance post-Bonferroni correction (*P*<0.05) and for suggestive significance (accounting for one false positive per genome scan).

**Fig 2.**
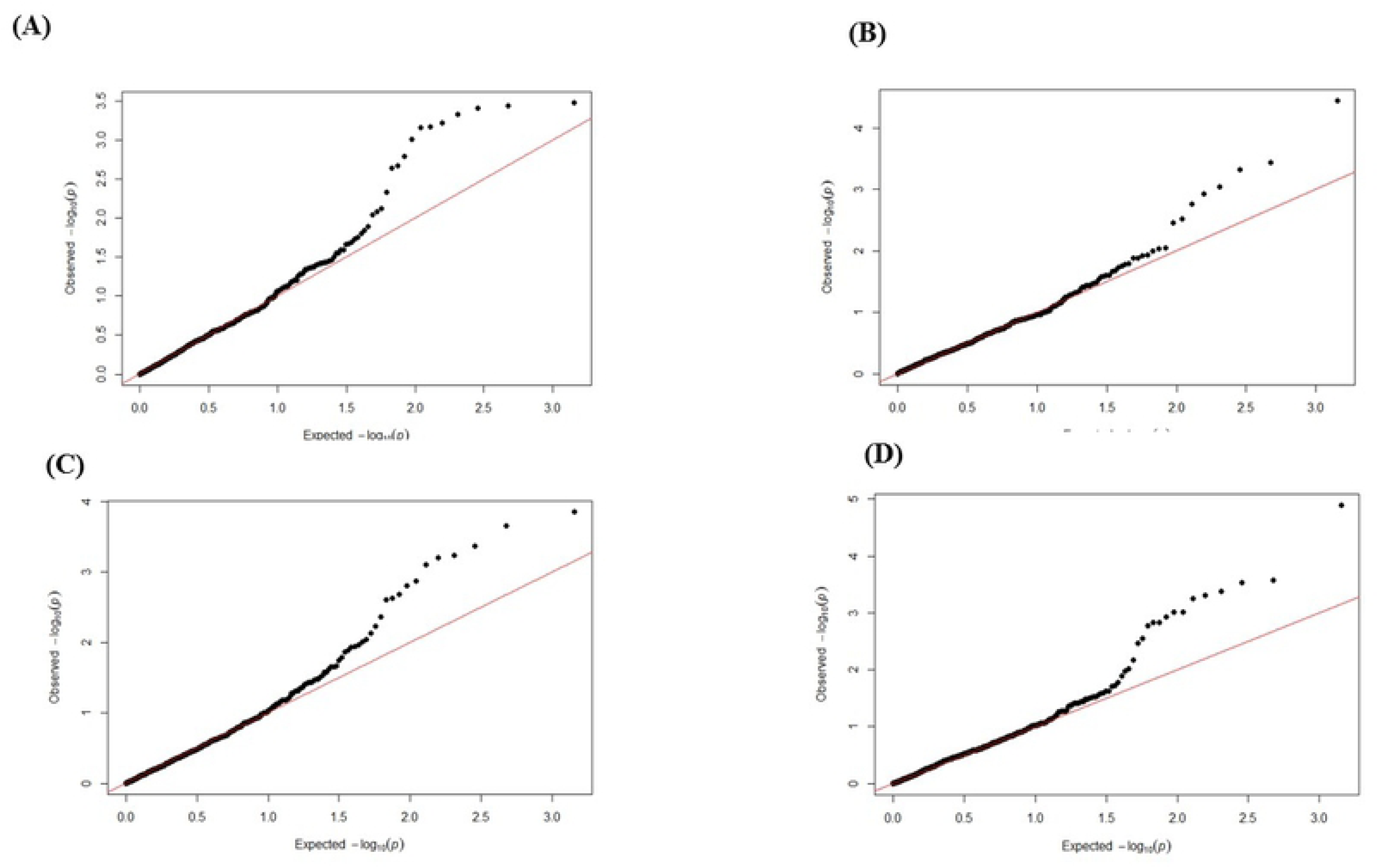
Q-Q plots displaying the genomic association results for milk yield in Chios sheep. Q-Q plots in early (A), mid (B), late (C) and overall (D) lactation; observed *P-*values are plotted against the expected *P*-values.

The significance of the above SNP markers was confirmed in mixed model analyses based on the pedigree genetic relationship matrix. The additive and dominance genetic effects, and the proportion of the total genetic variance explained by each of these SNPs in the corresponding lactation stage are summarised in Table 3. Most SNPs had a significant additive effect and a few a significant dominance effect on milk yield. The significant SNPs in the QTL region on chromosome 19 accounted for 16% of the additive genetic variance, while collectively all the SNPs in the four candidate regions accounted for 30% of the additive genetic variance of milk yield. When located on the same chromosomes, the significant markers identified for milk yield were in linkage disequilibrium (LD=0.27-0.97), implying that they correspond to the same causative mutation (S1 Table). The significant SNPs identified in the present study were not in LD with the SNPs previously associated with the mastitis related traits in Chios sheep [10] (S1 Table). Only small LD blocks were visualised with Haploview, probably due to a high number of recombination events having taken place in the outbred population of study. All significant SNP markers were located in intergenic or intronic regions. The candidate QTL regions for milk yield contained a relatively small number of protein-coding genes (n=31) and microRNAs (n=6) (S2 Table).

### Pathway and functional clustering analysis

The genes located in the candidate regions for milk yield were enriched for pathways involved in electrolyte (Na^+^, K^+^, and H^+^) transport and homeostasis, lipid metabolism (ketolysis, ketogenesis) and oxidative stress, as well as innate immune responses (Fig. 3). Moreover, two networks of molecular interactions were constructed, one of which was related with immunological disease and cell signalling and interaction, and another with the development, function and organ morphology of the endocrine and reproductive systems (Fig. 4).

**Fig 3.**
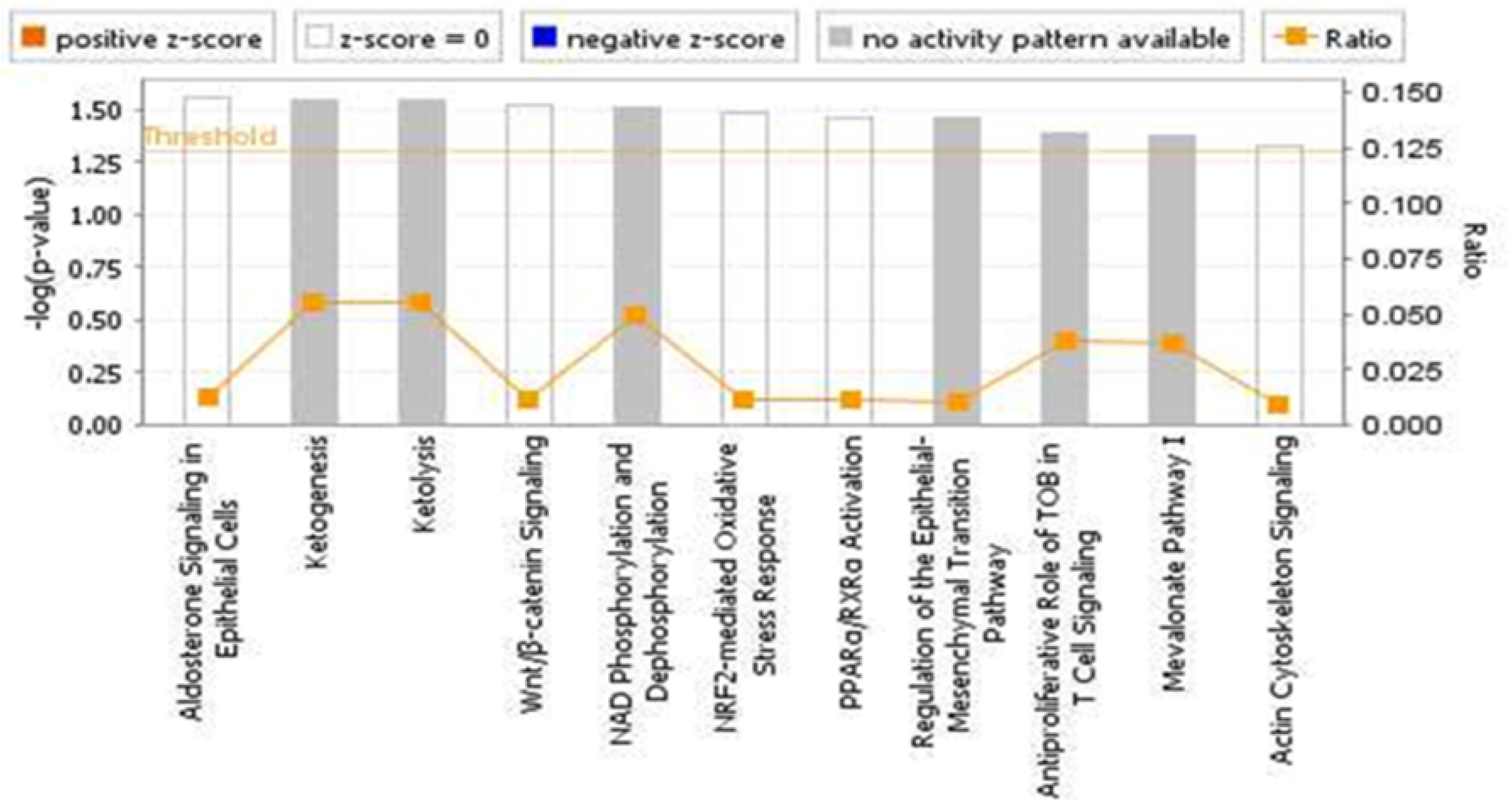
Pathway analysis using the IPA software. The most highly represented canonical pathways derived from genes located within the studied candidate regions for milk yield in Chios sheep. The solid yellow line represents the significance threshold. The line joining squares represents the ratio of the genes within each pathway to the total number of genes in the pathway.

**Fig 4.**
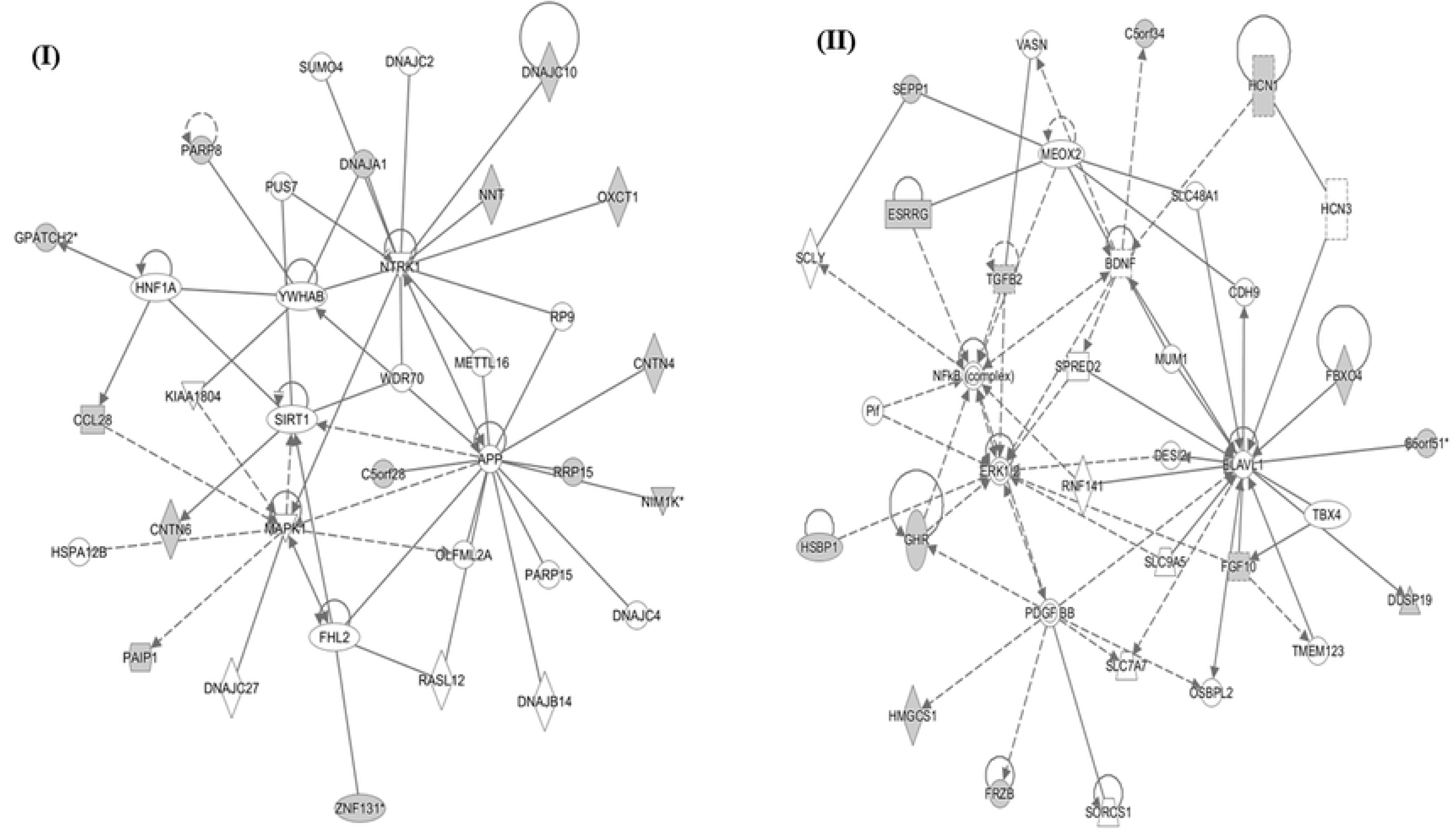
Network analysis using the IPA software. Two gene networks illustrating the molecular interactions between candidate gene products: (I) network related to immunological disease and cell signalling and interaction; (II) network related to the endocrine system development and function, reproductive system development and function, and organ morphology. Arrows with solid lines represent direct interactions and arrows with broken lines represent indirect interactions. Genes with white labels are those added to the IPA analysis because of their interaction with the target gene products.

The functional annotation clustering analysis showed that genes were organised into two clusters associated with the regulation of cellular processes (enrichment score =1.60) and metabolic processes (enrichment score =1.04). Both clusters contained the same genes (*CNTN4, DNAJA1, ESRRG, FGF10, FRZB, GHR, HMGCS1, OXCT1, TGFB2*).

### Gene expression analysis

Fourteen of the genes located in the candidate regions for milk yield (*CCL28, DNAJC10, DUSP19, EMB, FGF10, GHR, LYPLAL1, NNT, HMGCS1, NCKAP1, NUP35, OXCT1, PAIP1* and *ZNF131)* were expressed in either the milk transcriptome or the mammary gland (S1-3 Figs). The growth hormone receptor (*GHR*) and 3-oxoacid CoA transferase 1 (*OXCT1*) genes were highly expressed in liver and kidney cortex tissue, respectively (S2 Fig). Five of the candidate genes (*DNAJC10, EMB, HMGCS1, OXCT1, PAIP1*) detected in tissues related to milk production (mammary gland, liver and kidney cortex) were also up-regulated in immune related tissues, relative to the other tissues analysed (S3 Fig). The *EMB* gene in particular exhibited a strong immune-specific profile with a high level of expression in macrophages relative to the other tissues (S2 Fig).

Expression of genes *LYPLAL, PARP8, RRP15* and *TGFB2* had a suggestive (nominal *P<*0.05) association with milk yield in the Churra and Assaf sheep data that did not remain significant after the FDR correction.

A subset of five genes (*EMB, FGF10, DNAJC10, OXCT1, PARP8*) had a suggestive (nominal *P<*0.05) association with milk somatic cell count in the Churra and Assaf sheep data. The expression level of two of these genes, *EMB* and *FGF10,* were also significantly different between sheep with high and low somatic cell count in milk (Tukey’s Test; *EMB*: *P*=0.001648; *FGF10*: *P*=0.002085).

### Allelic expression imbalance analysis

Exonic single nucleotide variation (SNP and indels) was observed in 24 of the protein coding genes located in the candidate regions for milk yield. Missense variants were identified in several genes including, *CNTN4, DNAJC1, DUSP19, GHR, HMGS1, MRPS30, NNT, NUP35* and *RRP15* genes. One-sampled MBASED analysis identified two genes, *CCL28* (*P= 1.8e-05)* and *RRP15* (*P= 3e-03*) with significant allelic expression imbalance. Specifically, seven SNPs in the 3’ UTR region of *CCL28* (major allele frequency 0.72) and two synonymous SNPs in *RRP15* (major allele frequency 0.71) were detected exhibiting allelic expression imbalance (S3 Table). However, these results should be interpreted with caution since allelic expression imbalance in both genes was evident in only one of the three individual sheep.

### Candidate genes

Based on all above results, a total of seven genes (*DNAJA1, DNAJC10, FGF10, GHR, HMGCS1, LYPLA1, OXCT1*) were selected as candidate genes for milk yield located in mastitis genomic regions (Table 4). Genes were selected using a combination of their known biological function, involvement in relevant pathways and networks, enrichment in tissues relevant to milk production, and any previously known association with milk production in either dairy sheep or other species.

**Table 4.**
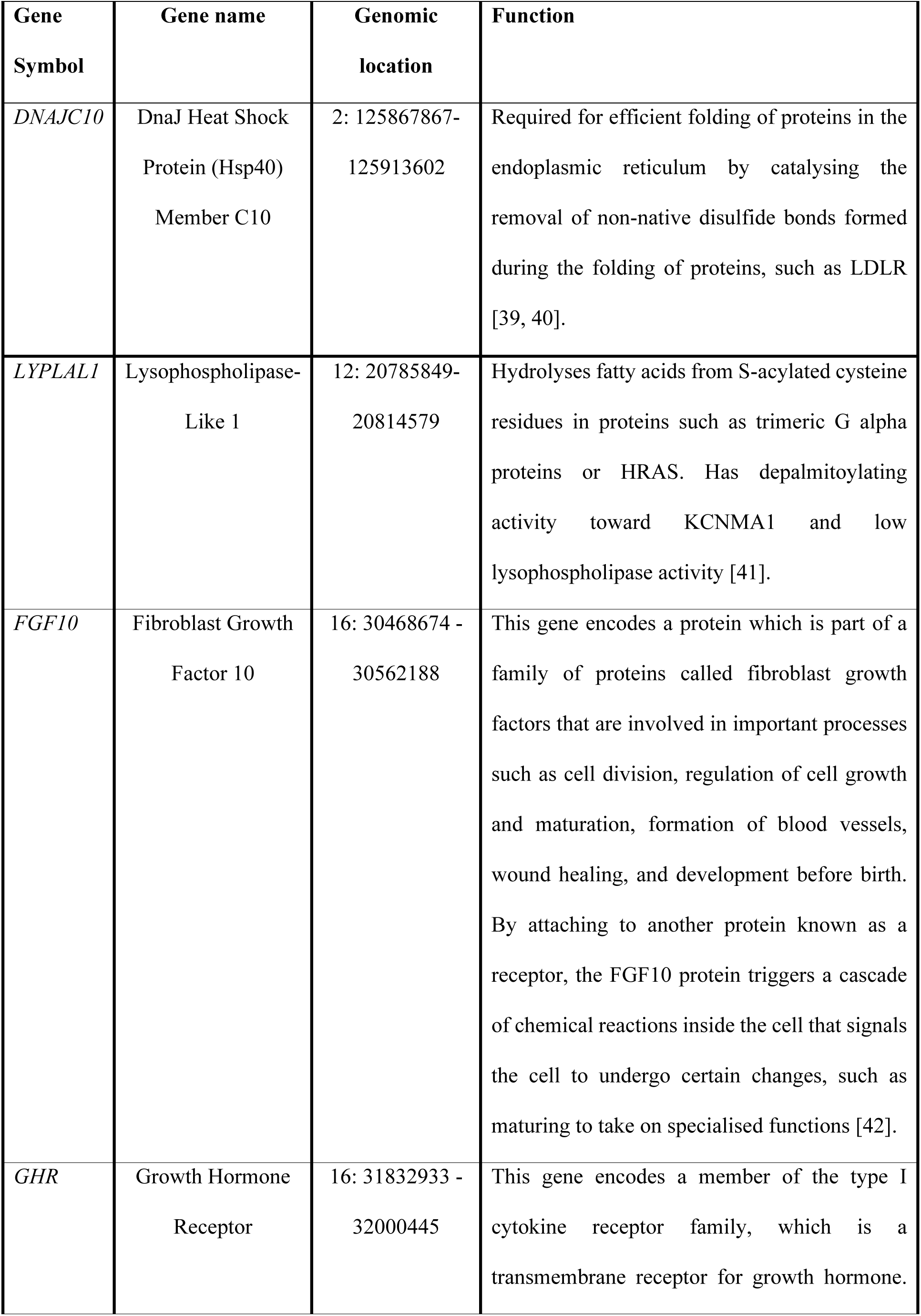
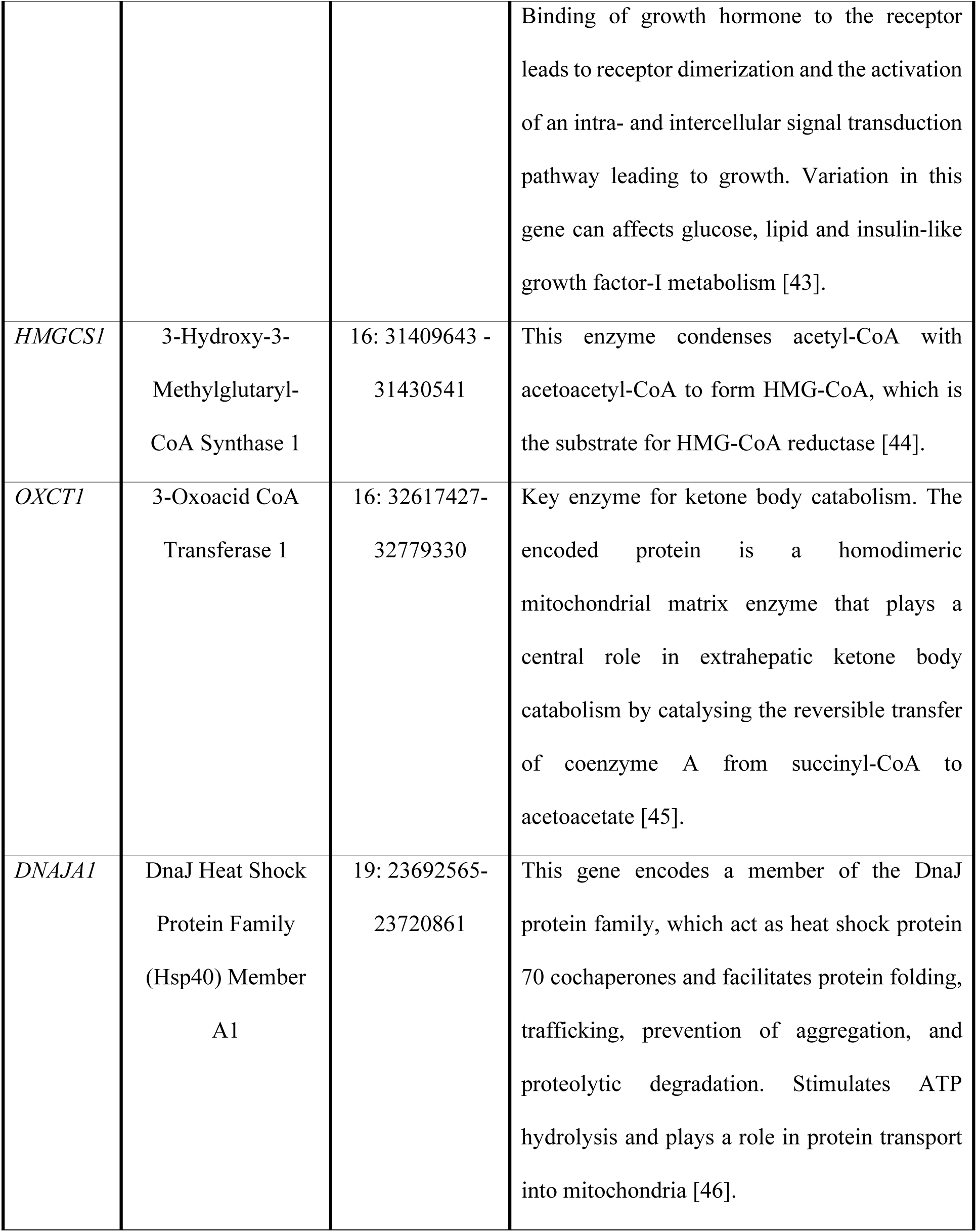
Selected candidate genes for milk yield.

## Discussion

The existence of a mastitis-specific ovine DNA array built on previously detected markers associated with mastitis resistance in dairy sheep opens up opportunities for targeted genomic and marker-assisted selection aiming to enhance animal resistance to the disease. The aim of the present study was to investigate the association of this array with milk yield of dairy sheep and assess the feasibility of a concomitant genetic improvement programme for the two traits. Chios sheep were used as a study model.

According to our results, milk yield and mastitis traits in the Chios sheep are not genetically correlated to each other. Genetic correlation estimates between milk somatic cell count and milk yield are reportedly antagonistic in dairy cattle [47] but inconsistent amongst previous sheep studies ranging from antagonistic [48] to favourable [3]. Our findings for the Chios sheep indicate that selection for enhanced mastitis resistance could be incorporated into the current genetic improvement programme without incurring adverse effects on milk yield.

An overall moderate but significant heritability for milk yield was estimated in Chios sheep, consistent with the dairy sheep literature (ranging from 0.16 to 0.30) as reviewed in [49] and previous studies in Chios sheep ranging from 0.21 to 0.29 [50].

Genomic analyses conducted here revealed several SNPs on the mastitis array with a significant effect on milk yield. These milk-associated SNPs were not in LD with genomic regions found previously to affect mastitis resistance in the same population [10]. For example, the QTL for milk yield on chromosome 2 was 75 Mb distant from the one previously identified for mastitis resistance on the same chromosome [10]. The association of this QTL region with milk yield is supported by results of a previous genomic selection mapping study that compared dairy with meat sheep breeds to identify genomic regions for milk traits under selection [51]. In that study a highly homozygous region was detected in both Chios and Churra sheep in close proximity with our QTL region on chromosome 2 [51]. Furthermore, the QTL for milk yield on chromosome 12 identified in the present study was located within a previously identified QTL region for milk yield in East Friesian X Dorset cross sheep [52]. No QTLs for mastitis resistance have been identified on this chromosome in Chios sheep [10]. The QTLs on chromosome 19 and 16 identified in the present study were also independent from those previously identified for mastitis resistance on the same chromosomes in the Chios sheep; the latter were located 2-4 Mb away and were in zero LD with the milk-associated region of the present study. QTLs for milk yield, and milk protein and fat content have been also identified on chromosome 16 in Churra sheep [53], in close proximity with the QTL identified here in Chios sheep. To the best of our knowledge, the QTL on chromosome 19 is reported here for the first time. These results are also consistent with a previous study of Chios sheep [54], suggesting that a relatively major locus might be involved in ovine milk production. The QTL identified on chromosome 19 in the present study explained 16% of the genetic variance. Furthermore, the significant SNP markers identified for milk yield in our study collectively explained over 30% of the genetic variance of the trait, suggesting that the mastitis-specific targeted array can also be used for genomic selection to enhance milk yield.

In the QTL regions identified for milk yield on chromosomes 2 and 19 there are two candidate genes, *DNAJA1* and *DNAJC10*, both belonging to the same gene family. In the previous milk transcriptome study of the Churra and Assaf breeds, two other related genes, *DNAJA4* and *DNAJB2,* were reported as functional candidates for milk yield [55]. The *DNAJ* family of proteins regulate ATP hydrolysis activity, and facilitate protein folding, trafficking, prevention of aggregation and proteolytic degradation; *DNAJA1* functions as a co-chaperone and protects cells against apoptosis in response to cellular stress [56], while *DNAJC10* promotes apoptosis in response to endoplasmic reticulum stress. Therefore, these two genes might affect milk yield through both metabolism and mammary apoptosis; the latter has been associated negatively with lactation persistency (daily milk yield decline in late lactation stages) in dairy species [57].

Some of the candidate genes for milk yield identified in the present study have been previously reported in dairy cattle. For example, 3-oxoacid CoA transferase 1 *(OXCT1*) has been associated favourably with both milk production [58] and mastitis resistance [59], and has been suggested to regulate mammary gland metabolism and milk synthesis during mastitis infection in dairy cattle [60]. Using the gene expression atlas for sheep and the milk transcriptome dataset, *OXCT1* was found to be expressed in both mammary gland and immune tissues, and highly expressed in the kidney cortex indicating that it may play a similar role in sheep. Growth hormone receptor (*GHR*) has been previously associated with increased milk yield and reduced milk somatic cell count in several dairy cattle studies [60–64]. Selective sweeps were also identified in the *GHR* region when dairy with beef cattle were compared [65]. In the present study, *GHR* was expressed in the mammary gland and the milk transcriptome, and was highly expressed in liver, relative to the other tissues sampled for the sheep gene expression atlas (http://biogps.org/sheepatlas). Furthermore, fibroblast growth factor 10 (*FGF10)* and 3-Hydroxy-3-Methylglutaryl-CoA Synthase 1 (*HMGSC1)* genes have been previously associated with milk production in dairy cattle [61, 66]. The pleiotropic growth factor *FGF10* is reportedly required for mammary gland development in mice [67]. In sheep, this gene was highly expressed in the mammary gland and female reproductive tissues including the uterus and placenta (http://biogps.org/sheepatlas). In the present study, *FGF10* was found to be differentially expressed in sheep with different milk somatic cell counts, suggesting a possible role in mastitis resistance. Our pathway analysis showed that the products of genes *GHR, FGF10, HMGSC1* are part of a network related with the development, function and organ morphology of the endocrine and reproductive systems. However, further studies are needed to confirm the relevance of this network with milk production and identify the causative genes and mutations.

Allelic expression imbalance was detected in the epithelia-associated chemokine (C-C motif) ligand 28 (*CCL28)* and ribosomal RNA processing 15 (*RRP15)* gene*s* in mammary gland tissue. The *CCL28* gene encodes a protein that has been previously reported to demonstrate direct antimicrobial activity against mastitis infection in dairy cattle [68] and in humans [69–71]. This gene is upregulated during pregnancy and lactation, and considered vital for the ability of IgA-producing B cells to migrate to the mammary tissue during lactation [71]. In the present study, *CCL28* was expressed highly in the milk somatic cells but not in immune tissues implying a protective role mainly in the mammary gland, as has been shown in humans [69, 70]. The *RRP15* gene plays a role in cell cycle, cell proliferation and apoptosis [72]. Both these genes are linked to protective immunity and, as such, are likely to be under strong selection pressure. The two variants detected in *PPR15* were synonymous SNPs and, therefore, less likely to be relevant to gene function. The seven variants in 3’ UTR of *CCL28* could be more relevant to the genetic/transcriptional control of *CCL28* expression, since the turnover of mRNA is mostly regulated by *cis-*acting elements located in the 3’UTR regions [73]. Therefore, these SNPs might affect the corresponding phenotypic trait in sheep, possibly by disrupting miRNA binding as in the myostatin example from Texel sheep described in [74]. Further analysis using the miRWalk database [75] predicted that one of the SNPs on chromosome 16 (31362143 bp) that exhibited allelic expression imbalance is located within a microRNA binding site, in the 3’ UTR of *CCL28*. Interestingly, the same microRNA (bta-mir-29e) in dairy cattle has been already reported to be differentially expressed in bovine mastitis caused by gram positive bacteria [76]. However, further studies are needed to confirm this, since allelic expression imbalance in the two genes was identified in only one of the three individual sheep studied, implying this might be simply due to individual specific variation in expression. A wider analysis across multiple tissues would also help to determine if the allelic expression imbalance observed in the present study is indeed specific to genes associated with the mammary gland. Further studies could also investigate the relevance of the SNPs exhibiting allelic imbalance to gene function and quantify allelic expression imbalance in a wider subset of animals, preferably including animals of the Chios breed.

In conclusion, results of the present study suggest that genetic selection for enhanced host resistance to mastitis will not antagonise milk yield in Chios sheep. Therefore, a genetic improvement programme for enhancing both mastitis resistance and milk production is feasible for this breed. In addition, there are QTLs within the mastitis specific DNA array that may be used to further increase milk production with genomic selection. Genes within genomic regions associated with ovine milk production exhibited tissue-specific expression patterns and pathways similar to those observed in cattle indicating that the underlying genetic mechanisms are likely to be, at least partially, conserved between the two species. Moreover, several candidate genes were highly expressed in immune tissues and in milk implying a favourable pleiotropic effect or likely role in milk production during udder infection. These genes are suitable candidates for further investigation to determine if they can be exploited in breeding programmes for concomitant improvement of milk production and mastitis resistance or as novel therapeutics.

## Availability of data and materials

All the raw gene expression data comprising the Texel and the Texel × Scottish Blackface gene expression atlas is available in the European Nucleotide Archive (ENA) under accession number PRJEB6169 and PRJEB19199, respectively. To supplement the data from these two transcriptomic atlas projects, we also obtained expression levels from a milk transcriptomic study of the milk somatic cells of two Spanish dairy sheep breeds, Churra and Assaf (Gene Expression Omnibus (GEO) database accession number GSE74825 and NCBI BioProject ID PRJNA301615).

## Authors’ contributions

GBa, AP and GA conceived and designed the genetic study of Chios sheep and secured substantial funding. AP, GBr and GBa performed data collection, phenotyping, DNA extractions and genetic parameter analysis. AP and GBa collated and edited the genotyping data and performed the genomic analysis. DAH and ELC conceived and designed the sheep gene expression atlas and DAH secured the substantial funding. ELC and SJB created the sheep gene expression atlas and analysed the gene expression data for the atlas dataset and the milk transcriptome. AP performed the pathway and the TFBS analyses. AP and SJB extracted and annotated the re-sequencing data of the HapMap sheep. PD performed the allelic expression imbalance analysis with input from AP, ELC, SJB and DAH. GBa, DH, GA, ELC and AP interpreted these results. GBa and AP wrote the manuscript. All other co-authors provided manuscript editing and feedback. All authors read and approved the final manuscript.

## Acknowledgements

The authors thank the cooperation of the commercial farmers who allowed access to their respective flocks.

## Supplementary files

**S1 Table**. Linkage disequilibrium (LD) estimates (expressed as r ^2^) for the significant SNP markers identified in the genomic association analyses of milk yield and mastitis resistance in Chios sheep.

**S2 Table.** Genes located in the candidate genomic regions identified for milk yield in Chios sheep.

**S3 Table.** Allelic expression imbalance analysis using the one-sampled MBASED method.

**S1 Fig**. Expression level of genes located in the milk yield candidate regions, as extracted from the Churra and Assaf sheep milk transcriptome analysis. Expression level is estimated as the mean number of transcripts per million of all (5) experimental replicates and is represented here as a z-score per individual animal.

**S2 Fig.** Expression level of genes located in the milk yield candidate regions, across all cell lines/tissues. Expression level is estimated as the mean number of transcripts per million (TPM) of all five (5) experimental replicates and is represented here as a z-score per cell line/tissue. Data is obtained from the sheep gene expression atlas which includes data from Texel X Scottish Blackface and Texel sheep.

**S3 Fig.** Expression level of genes, located in the milk yield candidate regions, across both mammary gland and immune cell lines/tissues. Expression level is estimated as the mean number of transcripts per million of all five (5) experimental replicates and is represented here as a z-score per cell line/tissue. Data is obtained from the sheep gene expression atlas which includes data from Texel X Scottish Blackface and Texel sheep.

